# CROCOpy - A Python toolbox for the analysis of CRitical Oscillations and COnnectivity

**DOI:** 10.64898/2026.02.17.706438

**Authors:** Vladislav Myrov, Felix Siebenhühner, Sheng H Wang, Gabriele Arnulfo, Joonas Juvonen, Monica Roascio, Gaia Burlando, Alina Suleimanova, Joona Repo, Wenya Liu, Satu Palva, J. Matias Palva

## Abstract

‘CROCOpy’ is a light-weighted toolbox for the assessment of neuronal oscillations, and multiple observables of functional connectivity (phase synchronization, amplitude coupling, and cross-frequency coupling) and critical dynamics (avalanches, long-range temporal correlations, bistability, and functional excitation-inhibition ratio). It was developed to simplify the analysis of continuous electrophysiological recordings and, in addition to metric computation, also includes methods for narrow-band filtering and statistical analysis. It is device-agnostic and supports both GPU and CPU computations. The toolbox also provides detailed tutorials.

## 1 Statement of need

The brain is a complex dynamical system, where neuronal interactions across spatiotemporal scales give rise to neural oscillations. These oscillations are related to each other by various forms of functional connectivity, and exhibit various forms of scale invariance such as avalanche dynamics, long-range temporal correlations (LRTCs), and bistability. Such concurrent and diverse activity patterns are hypothesized to emerge when a system is operating near a critical transition between order and disorder, a conceptual framework known as *brain criticality*.

Increasing evidence suggests that functional connectivity and critical dynamics are closely related to each other. Studying these together can therefore offer a holistic picture of neuronal dynamics. We here present ‘CROCOpy’, a unified and open-source Python toolbox, which combines implementations of multiple metrics used to analyze neuronal oscillations, their functional connectivity, and (critical) dynamics. The metrics it provides can be used for both basic research and as biomarkers in clinical contexts. The code is openly available on the GitHub (https://github.com/palvalab/crocopy)

## 2 State of the field

While several toolboxes exist that offer implementations of metrics for either criticality or connectivity, to our knowledge, none of these include the existing metrics used in studies of critical dynamics and connectivity altogether, focusing on specific methods such as DFA exponent or connectivity. Importantly, most existing toolboxes are not optimized for speed and do not utilize GPU which limits their applicability to large datasets.

## 3 Software Design

‘CROCOpy’ was designed to be as simple and flexible as possible and operates with basic N-dimensional arrays. Many operations can be indiscriminately run on either CPUs or GPUs, flexibly using either numpy or cupy under the hood. For most metrics, ‘CROCOpy’ supports GPU acceleration for efficient analysis of very large datasets. Additionally, ‘CROCOpy’ implements routines for narrow-band filtering and several commonly used statistical tests, with tutorials provided on GitHub.

Observables can be grouped in three categories (Table 1):

Brief descriptions of the methods can be found in the next section while detailed background, usage tips and discussion of how to interpret these metrics can be found in Tutorials.

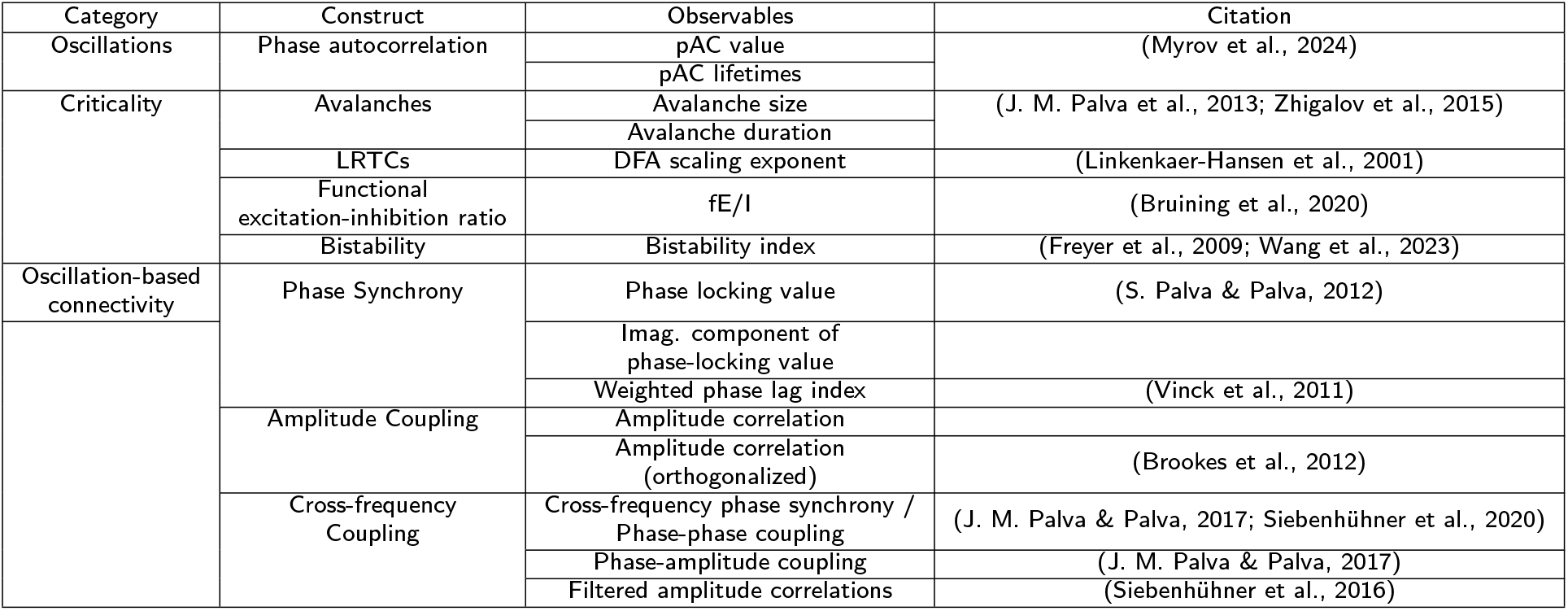

## 4 Implementation and core methods

### 4.1 Phase Autocorrelations

A neural oscillation “order” quantifies the local synchronization. Although its amplitude is a proxy for synchronization, it is not bounded to a specific interval and depends on other parameters. The phase autocorrelation function (pACF) quantifies the rhythmicity of a narrow-band signal by computing the self-similarity of its phase with a time-lagged version. The pACF operationalizes rhythmicity of oscillations either by the average value within a given lag range or by their lifetime (the lag at which the pACF crosses a specified threshold) (Myrov et al., 2024).

### 4.2 Criticality

In the *criticality* framework, the brain is hypothesized to be comparable to a system operating near a phase transition point, where its dynamics become scale-free and correlations extend across a wide range of temporal and spatial scales. Scale-free behavior indicates that a process has no single “typical” temporal or spatial scale that dominates its dynamics and is characterized by power-law scaling of relevant statistical quantities over orders of magnitude.

#### 4.2.1 Neuronal Avalanches

Neuronal avalanches in multichannel MEG/SEEG recordings are brief cascades of large-amplitude neuronal events that propagate across brain areas and exhibit power-law event size and lifetime distributions. ‘CROCOpy’ first detects in each channel the peaks (local maxima) whose amplitude exceeds a chosen threshold *T* (Figure 1A), convert each suprathreshold peak to a binary event, after which the time series is discretized into regularly-spaced time bins within which the number of spiking events are counted (Figure 1B). An avalanche is defined as a contiguous sequence of non-empty bins and its size is quantify as the total number of suprathreshold peaks in that sequence and its duration is the number of consecutive occupied bins (Zhigalov et al., 2015). Finally, the power-law is fitted to a distribution of avalanche sizes/lengths (Figure 1C).

**Figure 1:**
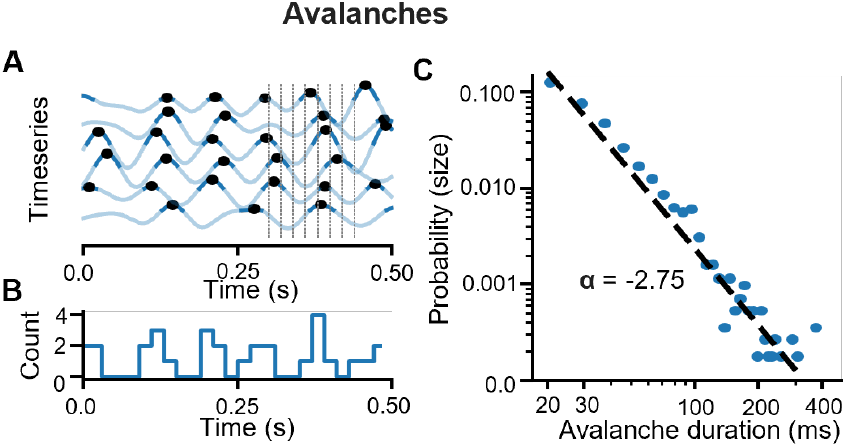
Avalanches. **A** Suprathreshold peak detection and time binning. **B** Peak counts per time bin. **C** Avalanche duration distribution with power-law fit.

#### 4.2.2 LRTCs

LRTCs in narrow-band oscillations describe how the amplitude of an oscillation fluctuates in a structured way over time. It can be assessed with detrended fluctuation analysis (DFA; Linkenkaer-Hansen et al., 2001). First, the signal profile is obtained as the cumulative sum of demeaned amplitudes, and detrended fluctuation analysis (DFA) is performed using logarithmically increasing window sizes (Figure 2A). For each window, a linear trend is fitted and removed, and the detrended fluctuation is quantified as the root-mean-square deviation from the fitted trend for that window size (Figure 2B). Finally, the DFA scaling exponent is obtained as the slope of a linear regression between log window size and log fluctuation. (figure 2C). This DFA procedure allows for the exclusion of “bad” samples (e.g., high-amplitude artifacts) but is slow to compute. We also implemented a spectral-domain approach to estimate signal variability (Nolte et al., 2019), which could be ten times faster but does not support artifact rejection.

**Figure 2:**
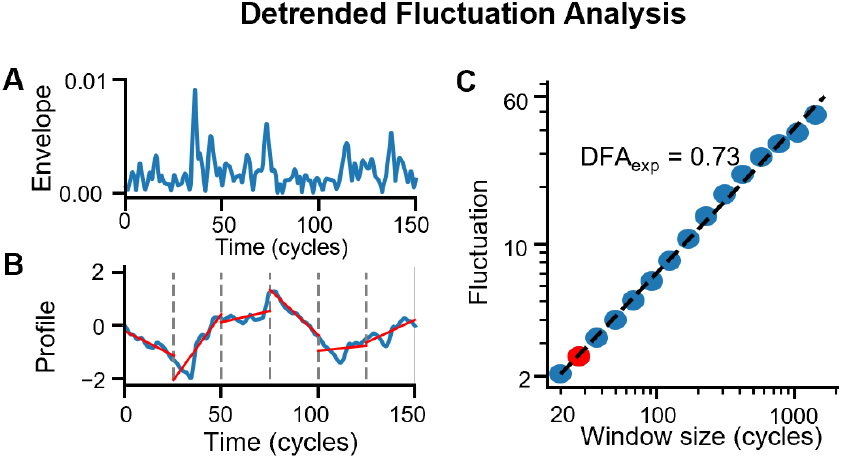
DFA. A Narrowband envelope time series. **B** Integrated profile with linear detrending in each window. **C** Fluctuation as a function of window size with linear fit (DFA exponent).

#### 4.2.3 Neuronal Bistability inside a Critical Regime

Bistability refers to dynamics that alternate between “down” and “up” states of synchrony, arising from positive feedback mechanisms, and is nonlinearly related to measures of criticality (Wang et al., 2023) (Figure 3A). First, narrow-band power time-series (squared envelope) is computed and normalized by maximum value (Figure 3B). Next, ‘CROCOpy’ compares whether its empirical distribution is better fitted by single exponential or bi-exponential function using Bayesian Information Criterion (BIC) (Figure 3C, (Freyer et al., 2009)).

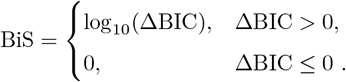

**Figure 3:**
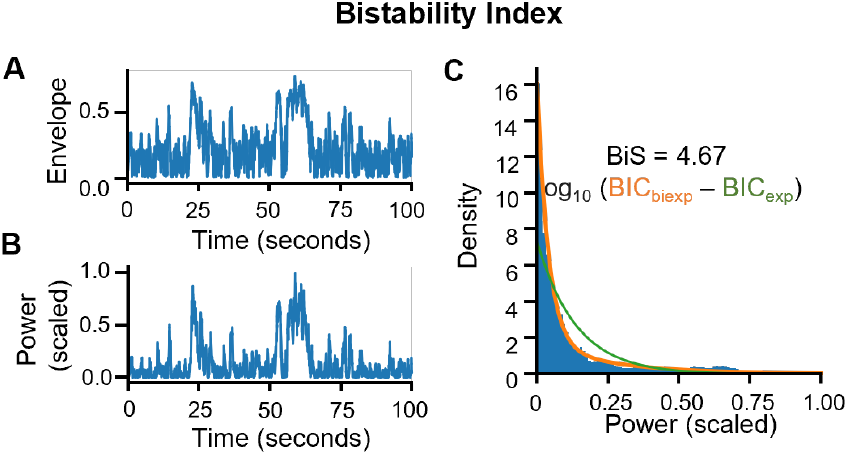
BiS. **A** Narrowband envelope time series. **B** Envelope power normalized to 0-1 range. **C** Power distribution with exponential and bi-exponential fits.

Thus, BIS > 0 indicates that the bi-exponential model (two-state structure) explains the power fluctuations better.

#### 4.2.4 Functional Excitation-Inhibition ratio (fE/I)

The fE/I index is intended to infer the local E/I ratio from a time series by quantifying how a critical oscillation (e.g., with DFA > 0.6) co-varies with the short-timescale temporal structure of amplitude fluctuations (Bruining et al., 2020). Like DFA analysis, the narrow-band signal profile is obtained (Figure 4A) and epoched into overlapping windows, and the average envelope is computed for each window (*W*_*env*_) (Figure 4B). Next, each window is amplitude-normalized by the corresponding mean envelope, detrended, and the fluctuation within the window is defined as the standard deviation (*W*_*fluct*_). The fE/I ratio is defined as *f E*/*I* = 1−*ρ*(*W*_*env*_, *W*_*fluct*_), where *ρ*(*W*_env_, *W*_fluct_) is the Pearson correlation across windows between their envelope and fluctuation (Figure 4C). The *f E*/*I* is bounded between 0 and 2: *f E*/*I* < 1 indicates inhibition-dominated (subcritical) dynamics, *f E*/*I* > 1 excitation-dominated (supercritical) dynamics, and *f E*/*I* ≈ 1 a near-critical E/I balanced regime.

**Figure 4:**
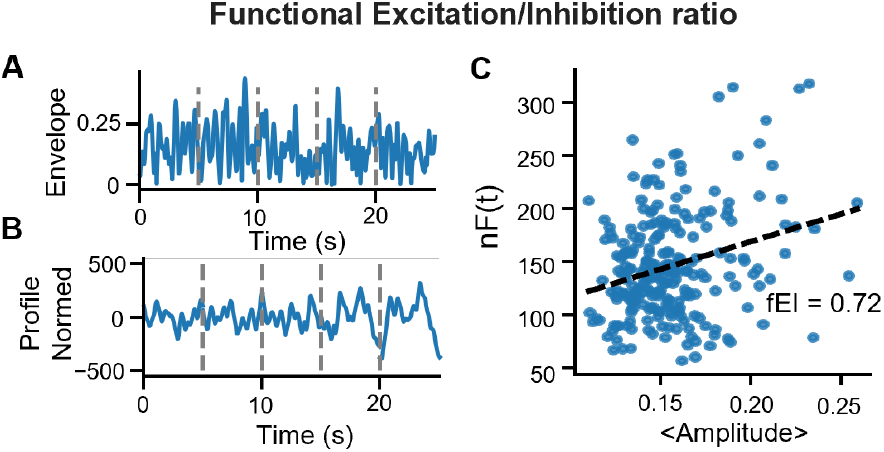
fE/I. **A** Narrowband envelope time series. **B** Integrated profile normalized by the window standard deviation. **C** Correlation across windows between mean amplitude and normalized fluctuation.

**Figure 5:**
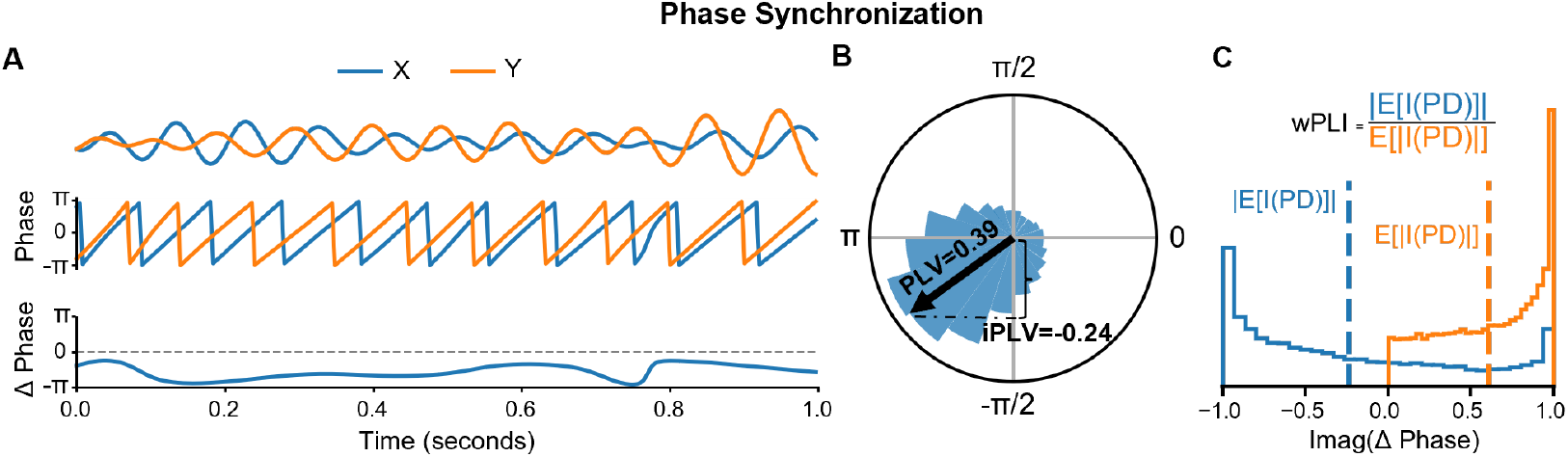
Phase synchronization. **A** Narrowband signals *X* and *Y*, their phases, and phase difference Δ*θ*(*t*). **B** Phase-difference distribution (circular plot) and corresponding imaginary-part distributions used for PLV/iPLV/wPLI.

### 4.3 Functional connectivity (FC)

FC denotes the statistical dependencies between neurophysiological signals recorded from different neuronal populations and is interpreted as inter-areal functional relationships that support communication and information transfer (Fries, 2015; S. Palva & Palva, 2012). FC in electrophysiological data comes in several types and can be estimated across a wide range of frequencies.

#### 4.3.1 Phase synchrony (PS)

PS refers to coupling between brain regions via a temporally stable phase difference between narrow-band signals (Figure 6A, (S. Palva & Palva, 2012)). With ‘CROCOpy’, PS can be estimated using the phase locking value (PLV), the imaginary phase-locking value (iPLV), and weighted Phase Lag Index (wPLI). EEG/MEG data are affected by linear signal mixing, which induces spurious interactions at zero phase lag (J. M. Palva et al., 2018). To avoid these false positives, iPLV only uses the imaginary component of the complex phase lag (which is zero when the phase lag is zero or a multiple of pi, Figure 6B), while the wPLI down-weights phase relationships near zero lag by weighting the phase difference by the magnitude of the imaginary component (Figure 6C).

**Figure 6:**
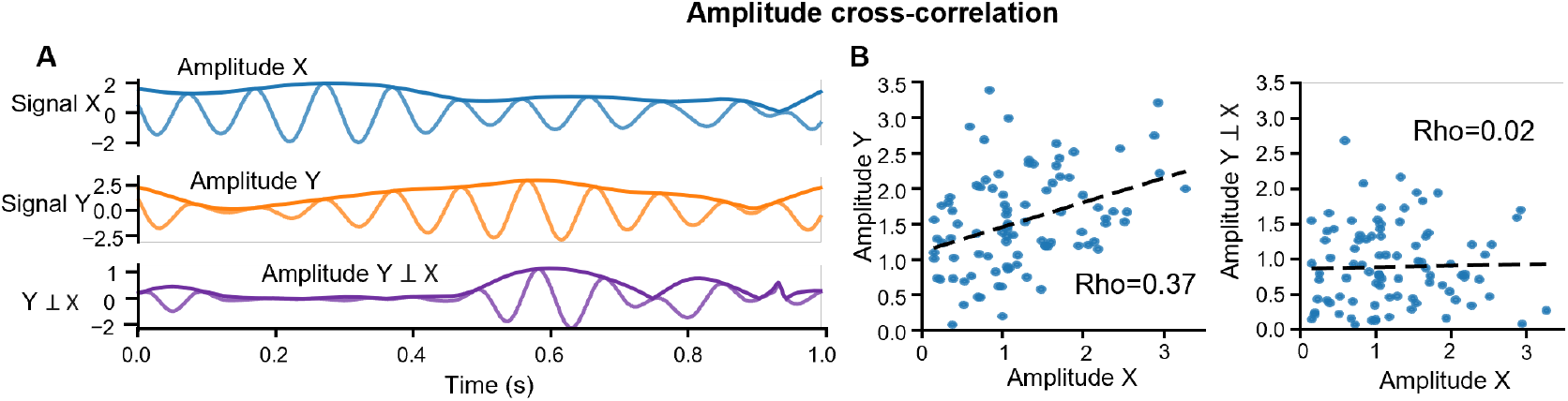
Amplitude correlations. **A** Narrowband amplitude signals and their envelopes *X* and *Y*, and *Y*_⊥*X*_ (orthogo-nalized w.r.t. *X*). **B** Amplitude correlation for *X* vs. *Y* (left) and *X* vs. *Y*_⊥*X*_ (right).

Both PLV and iPLV can be derived from the complex-valued PLV which is defined as:

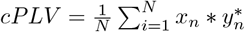

where *X*_n_ is unit-normed complex signal 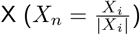 and 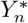 is conjugate of unit-normed complex signal Y. This yields a complex value, from which the regular PLV (|*cPLV* |) and the iPLV (|*img*(*cPLV*)|) can be obtained.

#### 4.3.2 Amplitude coupling (AC)

AC reflects inter-areal coupling of local fluctuations in neuronal firing patterns. Most simply, AC can be assessed by computing the Pearson correlation coefficient (CC) between envelopes of complex time series from two brain areas. However, CC is inflated by zero-lag artificial interactions as is the case for PS. To address this, the orthogonalized CC (oCC) was developed (Brookes et al., 2012), in which the correlation is computed between signal X and Y orthogonalized with respect to X (*Y* ⊥ *X*) as (Figure 6A,B):

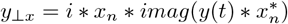

### 4.4 Cross-frequency coupling (CFC)

Various studies have indicated that neuronal oscillations interact not only within frequency bands, but also through multiple forms of cross-frequency coupling (CFC). These can be computed between different frequencies for the same region (local CFC) or between regions (long-range CFC) (Siebenhühner et al., 2020). Two widely investigated forms of CFC are cross-frequency phase synchrony (CFS) and phase-amplitude coupling (PAC).

CFS, also known as phase-phase coupling, is an extension of PLV to two oscillations with an integer frequency ratio n:m. CFS is estimated by estimating the phase-locking after phase-accelerating the slower oscillation (i.e., by multiplying its phase by m/n, Figure 7A). As CFC is assumed to be largely unaffected by linear mixing, it is not necessary to discard the real component of the cPLV (J. M. Palva et al., 2018).

**Figure 7:**
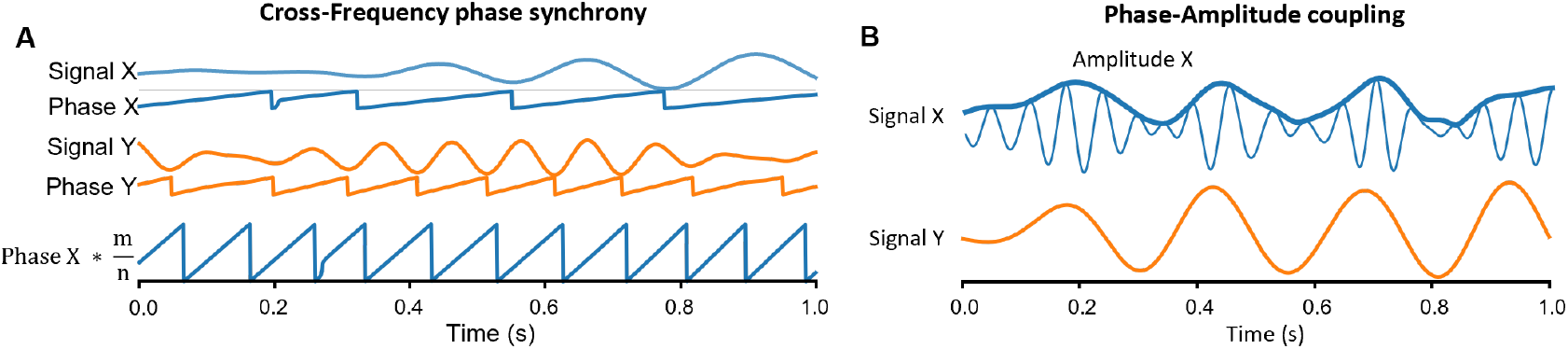
Cross-frequency coupling. **A** Narrowband signals *X* and *Y*, their phases, and the phase of *X* multiplied on 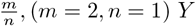, (*m* = 2, *n* = 1) *Y*. **B** Narrowband signals *X* and *Y* and the amplitude envelope of *X*.

PAC captures the dependence between the phase of a slower rhythm and the amplitude envelope of a faster rhythm, testing whether high-frequency power is systematically modulated at specific phases of the low-frequency oscillation. Several approaches exist for estimating PAC that generally yield comparable results. ‘CROCOpy’ implements PLV-PAC, which estimates PAC as the PLV between the slow oscillations’ phase and the phase of the amplitude envelope of the fast oscillation after wavelet convolution with the slow frequency (Figure 7B).

## 5 The critical and functional connectivity properties during sleep in pediatric cohort

Prior human studies have shown that sleep is accompanied by systematic changes in neural-dynamics observables—such as functional connectivity, long-range temporal correlations (LRTCs), and bistability—relative to wakefulness. Moreover, neural activity undergoes substantial reorganization across early neurodevelopment. However, the impact of early brain development on neural dynamics during sleep has not been systematically assessed. To provide an initial evaluation, we quantified functional connectivity and criticality-related metrics—DFA exponent, bistability index, and wPLI—using a previously collected pediatric sleep EEG dataset. Details of data acquisition and preprocessing are reported in the original data paper (Lee et al., 2022).

We found that in the youngest cohort (< 1 year) the bistability index has the lowest values, whereas in the oldest cohort (> 5 years) it reaches the highest values, with a peak in the beta (15–30 Hz) frequency range (Figure 8A). The DFA exponent in the youngest cohort (< 1 year) is also lower compared with the older groups and does not show a peak in the lower alpha band (6–10Hz) (Figure 8B). The connectivity profiles in the < 1 year group show two peaks, one around 4Hz and the other around 15Hz; with age, these peak frequencies move closer together and shift toward canonical alpha and beta bands in the > 5 years group (Figure 8C).

**Figure 8:**
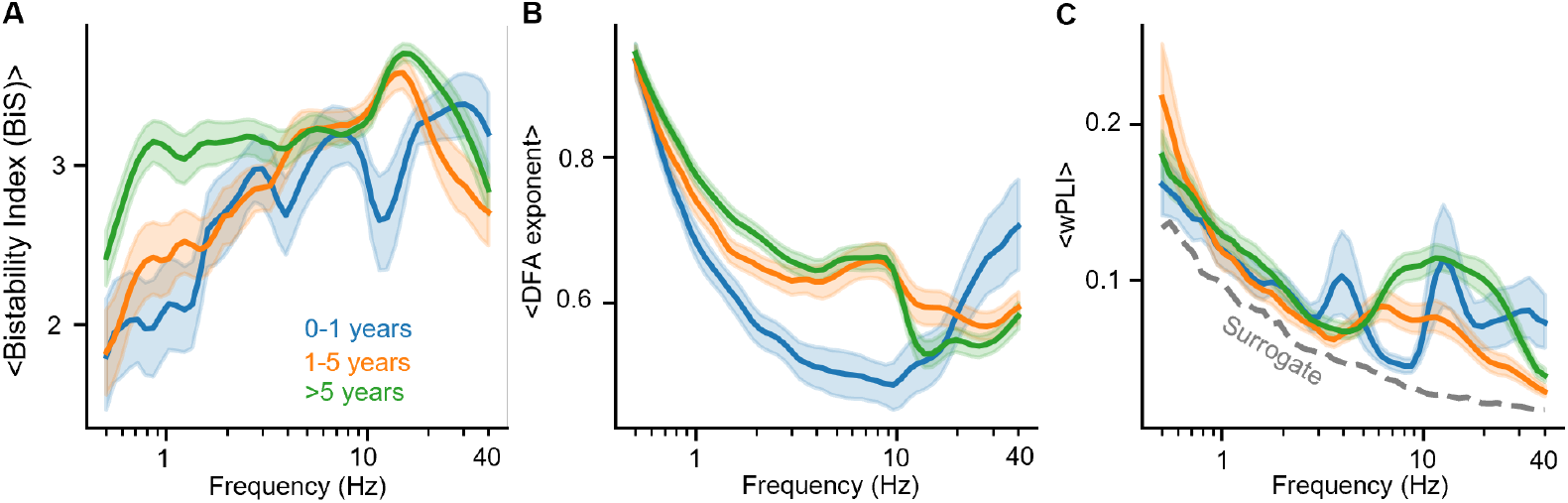
Critical and functional connectivity properties during sleep. An average Bistabilty index (**A**), DFA exponent (**B**) and wPLI (**C**) as a function of frequency for three cohorts: < 1 years old, 1-5 years old and *>* 5 years old. The shaded areas represent the variability computed from bootstraps (N=1000, 95th percentile).

## 6 Outlook and Future Efforts

The metrics described above reflect the implementation as of January 2026. We are a group of enthusiastic researchers actively developing novel criticality-inspired biomarkers, investigating their mechanistic links with established neurophysiological measures, and optimizing their use for studying complex neuroscience problems (Wang et al., 2025). Accordingly, this toolbox will be actively maintained, with ongoing improvements and the addition of new tools made available through GitHub. We warmly welcome contributions from researchers with shared interests and invite the community to participate in this long-term joint effort.

## 7 Research impact statement

CROCOpy has been used in multiple publications from several labs, including analyses of resting-state iSEEG functional connectivity of high-frequency oscillations, the relationship between DFA and phase synchronization, MEG/EEG data from patients with epilepsy, time-resolved features of multi-day ECoG recordings, and changes in brain dynamics during sleep.

## 8 AI usage disclosure

ChatGPT 5.2 was used for proof-reading the manuscript.

